# Boysenberry and apple juice concentrate reduces acute lung inflammation through increased alternatively activated macrophage activity in an acute mouse model of allergic airways disease

**DOI:** 10.1101/2020.03.03.974196

**Authors:** Odette M. Shaw, Roger D Hurst, Janine Cooney, Gregory M. Sawyer, Hannah Dinnan, Sheridan Martell

## Abstract

Bioactive compounds such as anthocyanins, proanthocyanins and other polyphenols are found in a wide variety of fruits and vegetables, and consumption of these have been associated with reduced lung inflammation and improved lung function in asthma and other lung diseases. We investigated whether a combination of Boysenberry and apple juice, found in BerriQi^®^ Boysenberry and apple juice concentrate, could reduce the allergic airways inflammation associated with asthma. We characterised the polyphenolic components in BerriQi^®^ Boysenberry and apple juice concentrate and identified the main compounds as cyanidin glycosides, ellagitannins, and chlorogenic acid. We found that consumption of 2.5 mg/kg of total anthocyanins from the BerriQi^®^ Boysenberry and apple juice concentrate significantly reduced eosinophil infiltration following acute ovalbumin (OVA) exposure in a mouse model of allergic airways inflammation. We found that BerriQi^®^ Boysenberry and apple juice concentrate consumption increased M2 (CD206+) macrophages and the production of the M2-associated cytokines CXCL10 and CCL4 within the lung. These results suggest that consumption of BerriQi^®^ Boysenberry and apple juice concentrate promotes a shift towards an anti-inflammatory environment within the lung leading to reduced immune cell infiltration and tissue damage.

## 1. Introduction

Asthma is a heterogeneous, chronic, inflammatory lung disease characterized by reversible airways obstruction, bronchospasm and infiltration of immune cells (1–3). It is estimated that 150 million people are affected by asthma worldwide, with a 5–15% prevalence in children (4) and there is evidence that early life exposure to air pollution caused by vehicle exhaust, environmental dust and industrial processes, increases the severity of asthma in children (5–7). The respiratory symptoms such as cough, and wheeze are worsened by exposure to pollution (8). Proinflammatory cytokine production in response to allergens by immune cells are further increased with concomitant pollution exposure (9–13). Eosinophils, in particular, produce reactive oxygen species, and cytokines, leading to epithelial damage and contribute to mucosal inflammation and the recruitment of other proinflammatory immune cells (14–17). These repeated acute inflammatory responses lead to tissue damage and remodelling, contributing to airway hyperresponsiveness, mucus cell hyperplasia, fixed airway flow obstruction, and loss of lung function over time (18–21).

Large-scale epidemiological studies have found that increased fruit and vegetable consumption correlates with reduced asthma symptoms (22–25). These dietary-related improvements in lung function benefits are also seen in people living in polluted environments (26–28). Fruits and vegetables contain numerous bioactive compounds, including anthocyanins and procyanidins, which have been shown to attenuate lung inflammation in cell and animal models of allergy and asthma (29–34). Human population studies have identified that dietary intake of foods high in polyphenols (28) such as apples, pears (35), carrots, tomatoes (25) and citrus is inversely correlated with the frequency and severity of reported asthma symptoms, especially wheezing and coughing (22, 25, 35, 36). Previously, we have identified that Boysenberry consumption led to decreased chronic lung inflammation, and improved lung tissue repair in an animal model of chronic allergic lung inflammation (33). Boysenberries contain high concentrations of anthocyanins and ellagitannins (37, 38). We have also found that procyanidin-rich extracts from apple suppressed IL-4 mediated cytokine production in cell culture models of lung epithelial allergic inflammation (32, 39).

There is increasing interest in understanding the mechanisms of action that specific plant bioactives have in the human body. This is partially to better understand the benefits of consuming specific fruits and vegetables, and partially to add value to specific foods through validated health claims. There is also interest in determining if combining specific plants containing different polyphenols can augment the health benefits above those seen with the individual plant. Use of animal models, where dietary intake can be tightly controlled are useful for both demonstrating/revealing the efficacy for identified compounds, and determining the biological mechanisms of action. The aim of this study was to determine whether the combination of Boysenberries and apple, as found in BerriQi^®^ Boysenberry and apple juice concentrate at a dose of 2.5 mg/kg total anthocyanins (TAC) could reduce allergic airways inflammation in response to acute ovalbumin exposure in a mouse model system. We also sought to determine the mechanisms involved in any ameliorating effect.

## 2. Methods

### 2.1 Mice and Materials

C57BL/6J male mice were group housed on 12h light/dark cycle in a conventional animal facility at The New Zealand Institute for Plant and Food Research Limited (Palmerston North, New Zealand). Mice were fed Prodiet RMH1800 standard chow for rodents (Lab Diet, St Louis, MO, USA) and filtered water ad libitum throughout the study, all attempts to minimise suffering were made. All experimental procedures were approved by the AgResearch Grasslands Animal Ethics Committee (AE approvals #14839, #14731 and #14016) and carried out in accordance with the Animal Welfare Act (1999). A proprietary Boysenberry and apple juice concentrate (BerriQi^®^) was supplied by Anagenix Ltd (Auckland, New Zealand). Legendplex^™^ 13-plex Th cytokine, proinflammatory cytokine and proinflammatory chemokine panels, Zombie NIR^™^ fixable viability dye, and anti-mouse CD3 (clone 17A2), CD4 (clone GK1.5), CD8a (clone 53-6.7), CD80 (clone 16-10A1), CD86 (clone GL-1), CD11c (clone N418), CD45 (clone 30-F11), CD206 (clone C068C2), CD14 (clone Sa14-2), Ly6C (clone HK1.4), Gr-1 (clone RB6-8C5), I-A/I-E (MHC class II; clone M5/114.15.2), and F4/80 (clone BM8) were purchased from Biolegend (San Diego, CA, USA). Anti-mouse SiglecF (clone E50-2440) and CD11b (clone M1/70) were from BD Biosciences (San Jose, CA, USA). Ovalbumin (OVA), and Alum were purchased from Sigma (Auckland, New Zealand). iScript Advanced cDNA kit was from Bio-Rad Laboratories (Hercules, CA, USA). Unless otherwise stated, all cell culture media, supplements, Taqman probes and buffers were purchased from Life Technologies NZ (Auckland, NZ).

### 2.2 BerriQi^®^ Formulation and Chemical Composition Analysis

Boysenberries (*Rubus ursinus* var loganbaccus cv Boysenberry), harvested between January-March 2017 from the Nelson region of NZ, were processed to a juice concentrate by Boysenberries NZ (Nelson, NZ). Apples (*Malus domestica* – mixed varieties) were harvested between April-June 2017 from the Hawkes Bay region of New Zealand, and juice concentrates prepared by Profruit Ltd. (Hastings, NZ). The freshly harvested Boysenberries and apples were graded, milled and pressed into juice, de-seeded by centrifugation, pasteurised, depectinised by enzymatic treatment, clarified by centrifugation, and concentrated by evaporation by their respective manufacturers to yield fruit juice concentrates, each with an 8-fold concentration factor.

Shott NZ Ltd. (Wellington, NZ) then blended the juice concentrates (proprietary proportions), and added potassium sorbate preservative to a final concentration of 0.12% (w/w), yielding a dark red, non-cloudy syrup. The final BerriQi^®^ Boysenberry and apple juice concentrate product measured pH 3.39 (by pH meter) and was 68.3° Brix (by refractometry), with a specific gravity of 1.35 mg/mL comprising 68% solids measured by both gravimetric and drying methods.

The polyphenol content of the BerriQi^®^ Boysenberry and apple juice concentrate was determined by liquid chromatography-mass spectrometry (LC-MS) using an LTQ linear ion trap mass spectrometer fitted with an ESI interface (ThermoFisher Scientific, San Jose, CA, USA) coupled to an Ultimate 3000 UHPLC and PDA detector (Dionex, Sunnyvale, CA, USA). Anthocyanins were separated on a Poroshell 120 SB-C18, 2.1×150 mm, 2.7 μm, analytical LC column (Agilent, Torrance, CA, USA), maintained at 70°C. The solvents were (A) 5:3:92 acetonitrile:formic acid:water v/v/v and (B) acetonitrile + 0.1% formic acid (flow rate, 200 μL/min). The initial mobile phase, 100% A, was held for 2 min before being ramped linearly to 88% A at 14 min, returning to 5% A at 15 min and held for 4 min before resetting to the original conditions. The sample injection volume was 10 μL. The MS data were acquired in the positive mode. Anthocyanin concentrations are reported as cyanidin-3-*O*-glucoside equivalents.

Other phenolic compound separation was achieved using a Hypersil GOLD aQ 1.9μ C18 175Å (Thermo Scientific, Waltham, MA, USA), 150 × 2.1 mm column maintained at 45°C. The solvents were (A) water + 0.1% formic acid and (B) acetonitrile + 0.1% formic acid (flow rate, 200 μL/min). The initial mobile phase, 95% A/5% B, was ramped linearly to 85% A at 10 min, held for 3.75 min, then ramped linearly to 75% A at 18 min, 67.2% A at 25 min, 50% A at 28 min, 3% A at 29 min and held for 4 min before resetting to the original conditions. The sample injection volume was 4 μL. The MS data were acquired in the negative mode.

### 2.3 OVA-Induced Airway Inflammation Model

Allergic airway disease was induced as previously described (34, 40). For the Boysenberry and apple interventions, mice were randomized into receiving either water (vehicle control) or 2.5 mg/kg TAC in the BerriQi^®^ Boysenberry and apple juice concentrate as previously described (34). Briefly, mice were fasted for 4 h before being orally gavaged with water (control) or at a dose of 2.5 mg/kg body weight TAC in the BerriQi^®^ Boysenberry and apple juice concentrate made up to a total volume of 200 μL in water 1h before OVA challenge and again 2 days post-challenge. Mice were sacrificed 4 days following intranasal ovalbumin challenge and immune parameters were analyzed.

### 2.4 Immune Parameter Analysis

Bronchoalveolar lavage fluid (BALF) and lung tissues were collected as previously described and immune cells were phenotyped by flow cytometry (40). Lung tissue supernatant for cytokine analysis was prepared as previously described (34). Briefly, the left lung lobe was minced into 500 μL phenol red-free complete RPMI media and incubated at 37°C for 30 min before being filtered through a 40 μm mesh and centrifuged to remove cellular material. The resulting supernatant was analysed for cytokine concentration using Legendplex bead-based multiplex immunoassays as per the manufacturer’s instruction. Both cell phenotyping and the cytokine multiplex assays were analyzed using a BD FACSverse (BD Biosciences, San Jose, CA, USA). H&E and AB-PAS histological staining were performed by Massey IVABS histology unit.

### 2.5 Real-time qPCR analysis

Mouse lung tissue was collected 4 days following OVA challenge and snap frozen in liquid nitrogen. The lung samples were crushed into powder using a mortar and pestle with liquid nitrogen to keep the samples frozen. The RNA was extracted from the powder using a TRIZOL total RNA extraction protocol. RNA was quantified using an LVis plate in a POLARstar Omega plate reader (BMG) and the quality of the ribosomal RNA bands confirmed by agarose gel electrophoresis (data not shown). Five μg of RNA from each sample was used as the template for cDNA synthesis using the iScript^™^ cDNA Synthesis Kit. Taqman^®^ Gene Expression Assays were purchased for each gene of interest. Two housekeeping genes, GAPDH and β-actin, were used as controls to determine the amount of relative gene expression. Taqman^®^ Gene Expression Master Mix was used to PCR amplify the genes in a Bio-Rad^™^ CFX384^™^ Real-Time PCR Detection System. Three lung samples per treatment group were prepared and amplified in quadruplicate with the housekeeping genes amplified on the same 384-well plate.

### 2.6 Statistical Analysis

Data were analyzed using one-way analysis of variance (ANOVA) with a Tukey’s post hoc test and graphed in SigmaPlot 12.5 (Systat Software Inc., San Jose, CA, USA).

## 3. Results

### 3.1 Chemical Composition of the Boysenberry and Apple Juice Concentrate

The results of the LC-MS analysis showed that cyanidin glycosides, ellagitannins, and chlorogenic acid were the major components in BerriQi^®^ Boysenberry and apple juice concentrate (Table 1, supplementary figures 1, 2). Minor components included phloretin 2-O-glucoside, and a mix of phenolic acids, flavonol glycosides, flavanol monomers and procyanidins. The major classes of phenolic compounds were anthocyanins (1969 μg/mL) and hydrolysable tannins (946 μg/mL), accounting for 56% and 27%, respectively, of the total phenolics quantified. The most abundant tannins were ellagic acid (449 μg/mL) and sanguiin H6 (213 μg/mL).

**Table 1.**
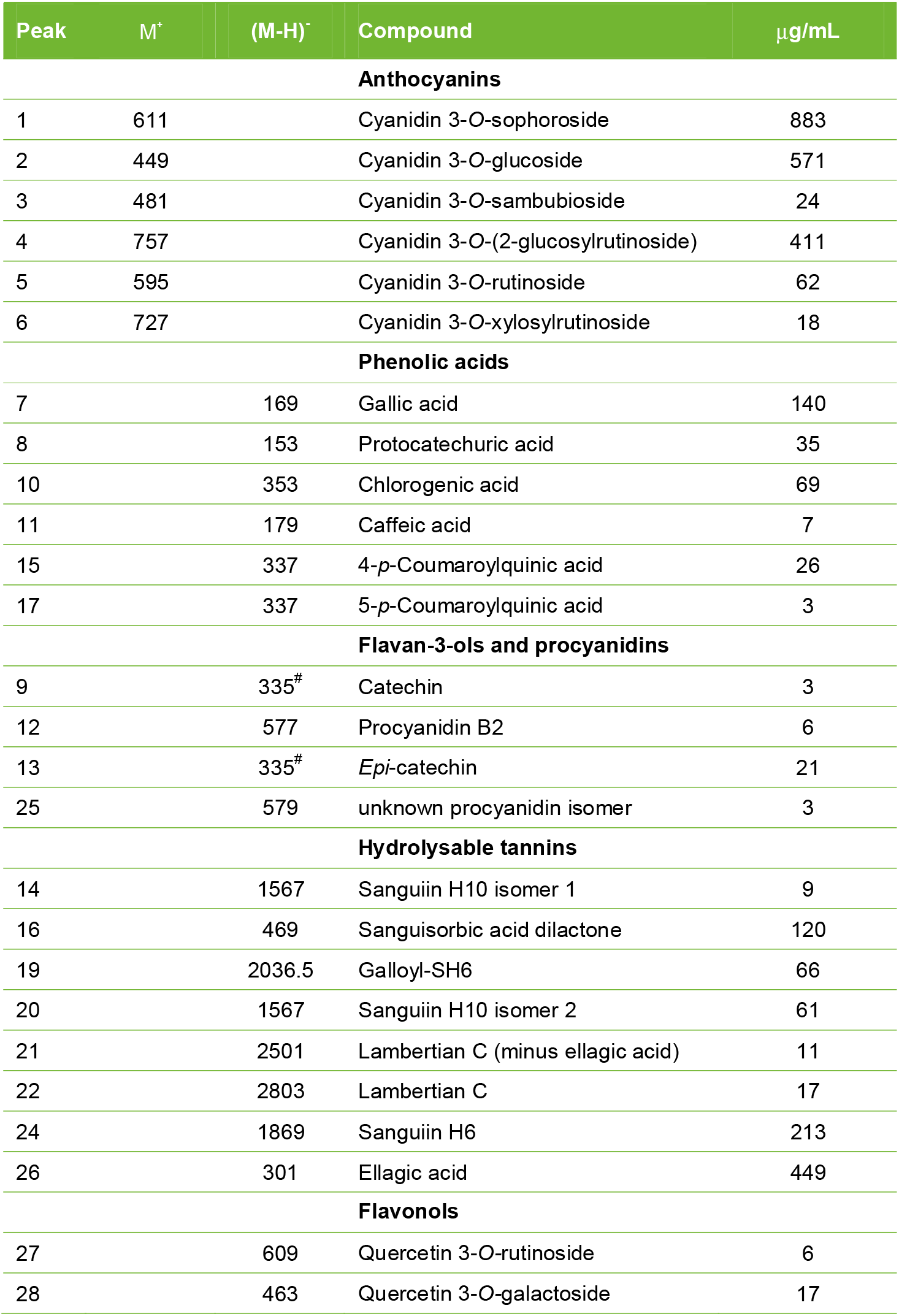
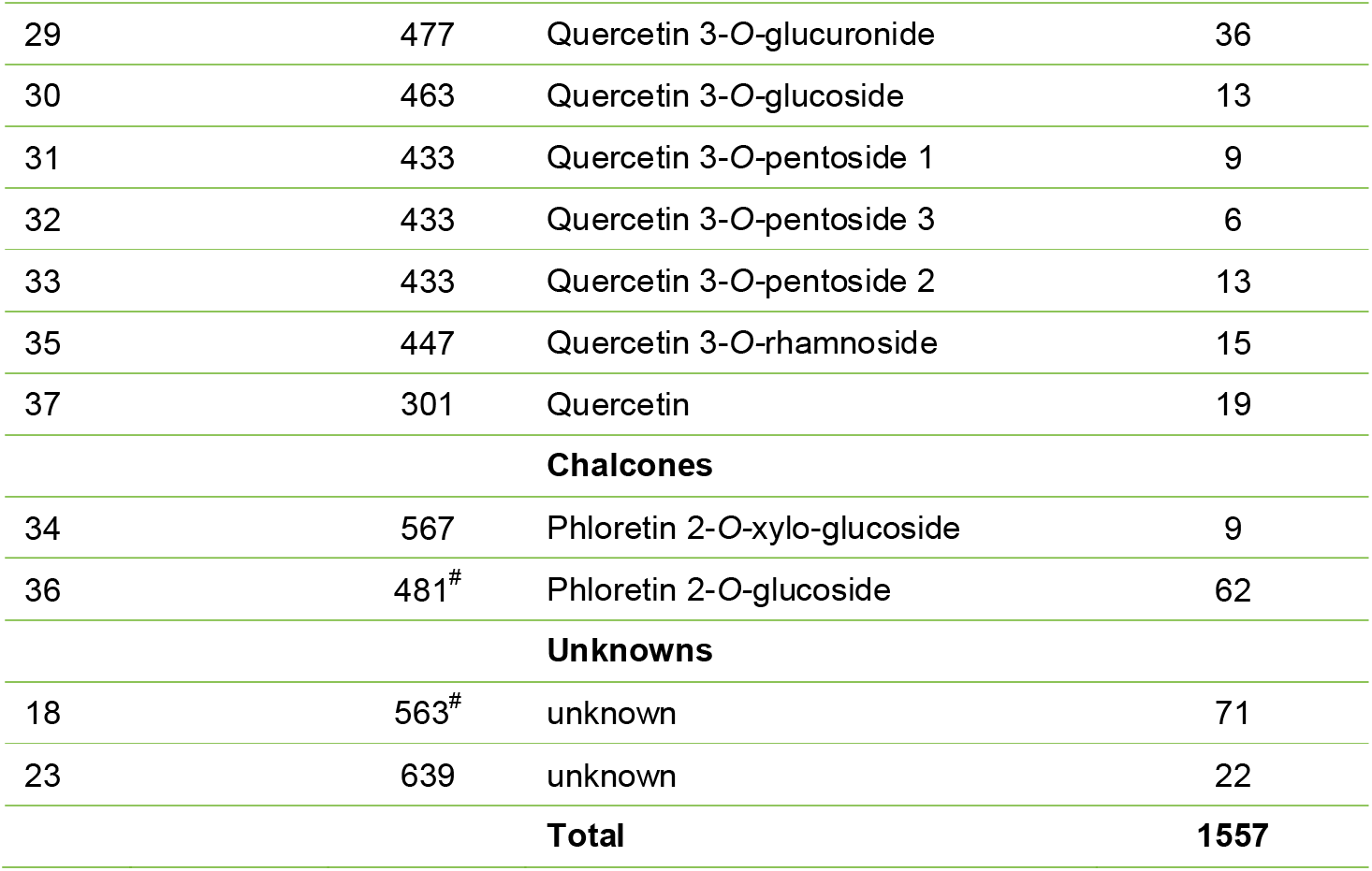
Phenolic compounds detected in BerriQi^®^ (μg/mL). M^+^ and (M-H)^−^ ions are the pseudomolecular ions used for identification of compounds by liquid chromatography-mass spectrometry (LC-MS). All identifications confirmed by MS/MS^n^ experiments. Peak numbers refer to chromatograms shown in supplementary data. # detected as [M+formate]^−^ adduct

### 3.2 Effect of Boysenberry and Apple juice concentrate Intervention on Ovalbumin-Induced Allergic Airways Inflammation

Acute intranasal OVA exposure resulted in an infiltration of immune cells into the lung (Figure 1A) and increased mucous production (Figure 1B). Consumption of 2.5 mg/kg TAC BerriQi^®^ Boysenberry and apple juice concentrate reduced the infiltration of immune cells and decreased OVA-induced mucous production (Figure 1A-B). We quantified the type and number of immune cells infiltrating into the lung, and found that acute OVA exposure significantly increased (P<0.001) infiltrating eosinophils (CD45+/CD11b+/SiglecF+), neutrophils (CD45+Ly6C+Gr-1+) and T-cells (CD45+/CD3+/CD4+ or CD45+/CD3+/CD8a+), compared with the lung of naïve animals (Figure 1C-F). Compared with animals only exposed to OVA, those that also consumed 2.5 mg/kg TAC BerriQi^®^ Boysenberry and apple juice concentrate showed a significant decrease (P<0.001) in the number of infiltrating eosinophils, neutrophils and T cells in the lung (Figure 1C-F). We saw no change in the number of CD4+ or CD8+ T cells in the mediastinal (lung-draining) lymph node for any of the treatment groups (Figure 2A-B).

**Figure 1:**
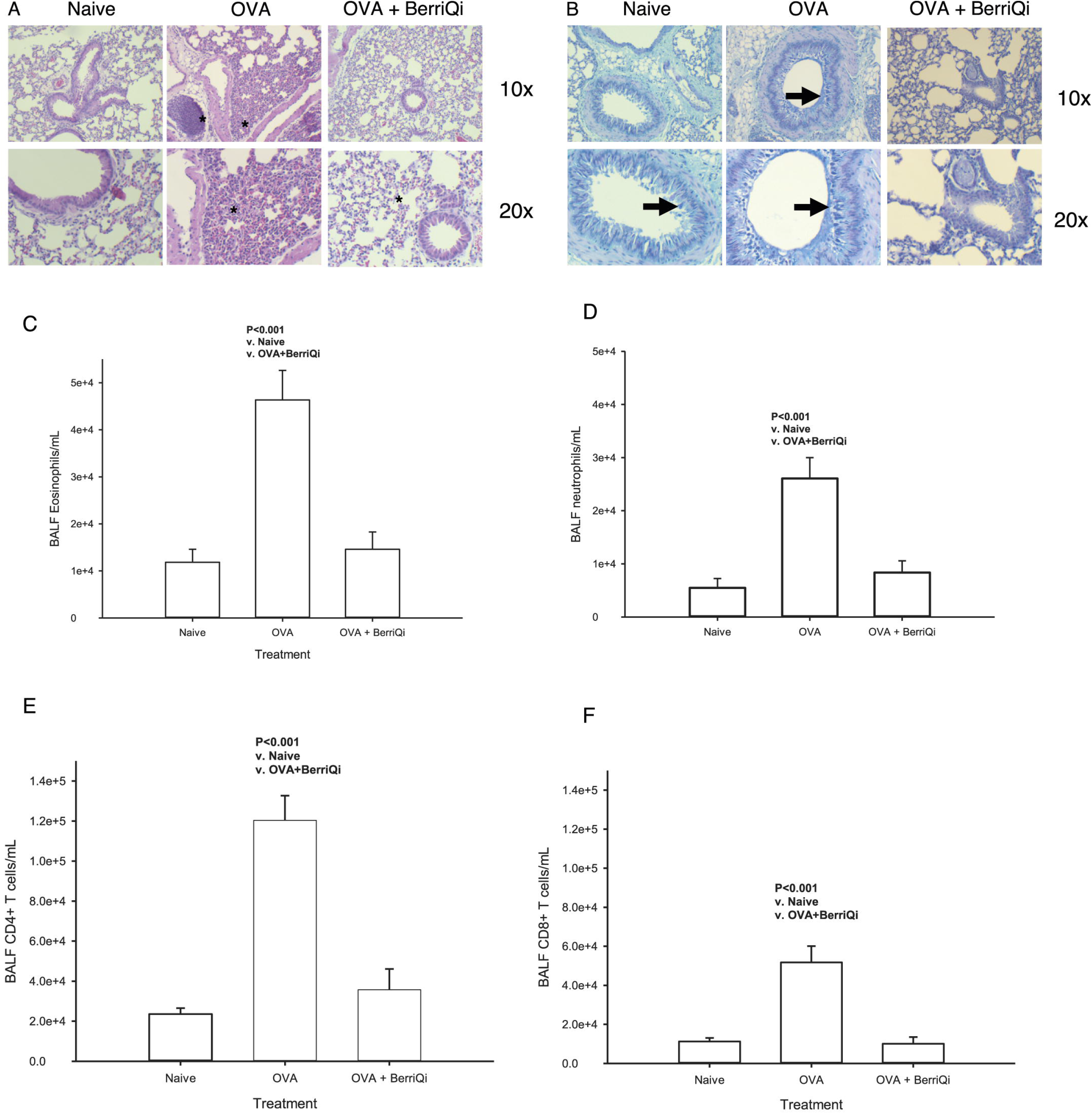
BerriQi^®^ Boysenberry and apple juice concentrate suppresses ovalbumin-induced airway inflammation, and immune cell infiltration. Mice were primed with ovalbumin (OVA)/Alum i.p and then challenged 7 days later with OVA i.n (Day 0). Mice were orally gavaged with 2.5 mg/kg total anthocyanins (TAC) in the BerriQi^®^ Boysenberry and apple juice concentrate (BerriQi) 1h before OVA challenge and again 2 days post-challenge. (A) Haematoxylin and eosin stained lung tissue from naïve, OVA-challenged and OVA-challenged mice treated with BerriQi^®^ Boysenberry and apple juice concentrate. Magnification 10x (top) and 20x (bottom) Asterisk =cell infiltration. (B) Alcian-blue Periodic acid-Schiff stained lung tissue from naïve, OVA-challenged and OVA-challenged mice treated with BerriQi^®^ Boysenberry and apple juice concentrate. Magnification 10x (top) and 20x (bottom). Arrow=mucous producing goblet cells. (C) Total eosinophil, (D) Total neutrophil, (E) CD4+ T cells and CD8+ T cells in bronchioalveolar lavage fluid (BALF) were determined 4 days post-OVA challenge. Data presented as mean ± SEM P<0.001 compared with naïve and OVA challenge + BerriQi^®^ Boysenberry and apple juice concentrate (one-way ANOVA with Tukey’s Post Hoc test) for two experimental replicates with n=10 per treatment groups.

**Figure 2:**
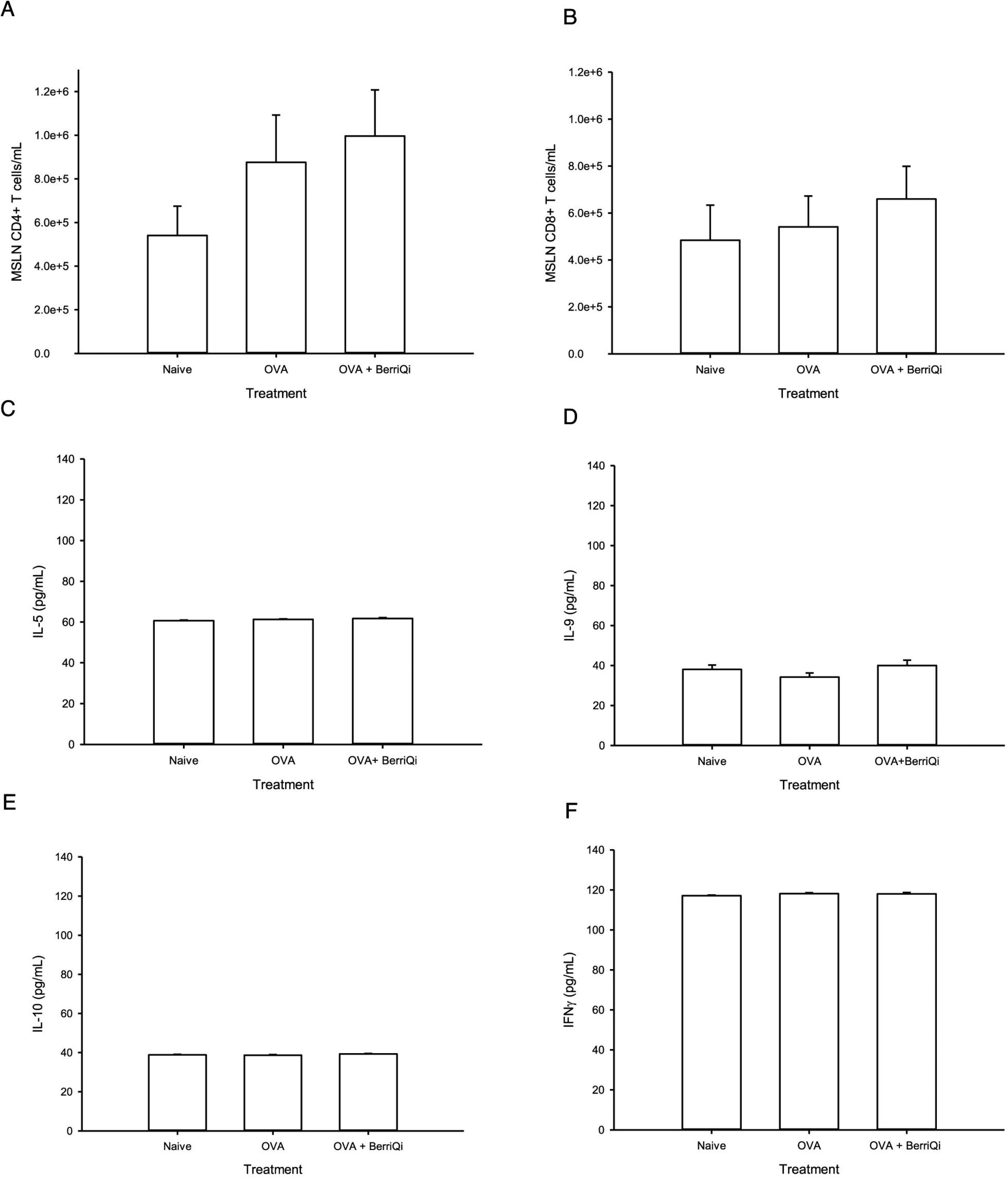
BerriQi^®^ Boysenberry and apple juice concentrate does not alter classical Th-1/Th-2 cells and cytokines. Mice were primed with ovalbumin (OVA)/Alum i.p and then challenged 7 days later with OVA i.n (Day 0). Mice were orally gavaged with 2.5 mg/kg total anthocyanins (TAC) in the BerriQi^®^ Boysenberry and apple juice concentrate (BerriQi) 1h before OVA challenge and again 2 days post-challenge. Mediastinal lymph node (MSLN) (A) CD4+ and (B) CD8+ T cells number; and lung tissue production of (C) IL-5, (D) IL-9, (E) IL-10 and (F) IFNγ were determined 4 days post-OVA challenge by Legendplex. Data presented as mean ± SEM for two experimental replicates with n=10 per treatment groups.

There was a trend towards an increased percentage of CD206+/CD14-macrophages in the lungs of mice that consumed 2.5 mg/kg TAC BerriQi^®^ Boysenberry and apple juice concentrate (Figure 3A). We measured the gene expression of Arg1, Ym-1 and Fizz1 in lung tissue and found that 2.5 mg/kg TAC BerriQi^®^ Boysenberry and apple juice concentrate consumption led to a significant fold-increase in Arg1 (P<0.05) and Fizz1 (P<0.01) gene expression compared to naïve mice (Table 2). We found no significant fold-change in Nos2 or Ym-1 gene expression between any of the treatment groups (Table 2).

**Figure 3:**
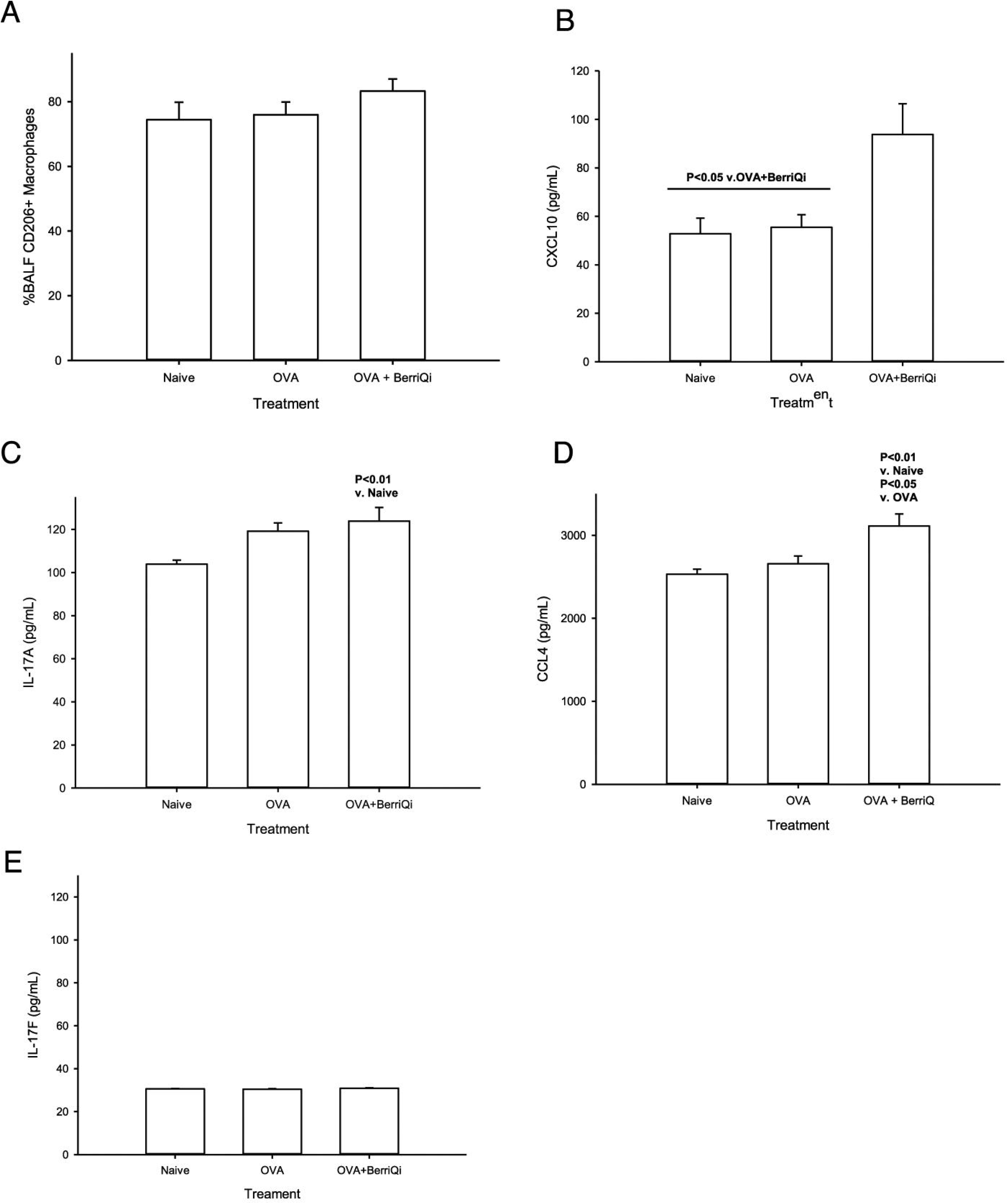
BerriQi^®^ Boysenberry and apple juice concentrate suppresses ovalbumin-induced airway inflammation through increased IL-17A, CXCL10 and CCL4 concentration. Mice were primed with ovalbumin (OVA)/Alum i.p and then challenged 7 days later with OVA i.n (Day 0). Mice were orally gavaged with 2.5 mg/kg total anthocyanins (TAC) in the BerriQi^®^ Boysenberry and apple juice concentrate (BerriQi) 1h before OVA challenge and again 2 days post-challenge. (A) Percentage of CD206+ macrophages in bronchioalveolar lavage fluid (BALF) and lung tissue production of (B) CXCL10, (C) IL-17A, (D) CCL4 and (E) IL-17F was determined 4 days post-OVA challenge by Legendplex. Data presented as mean ± SEM, P<0.05 compared with OVA challenge + BerriQi^®^ Boysenberry and apple juice concentrate, P<0.01 compared with naïve and OVA challenge (one-way ANOVA with Tukey’s Post Hoc test) for two experimental replicates with n=10 per treatment groups.

**Table 2:**
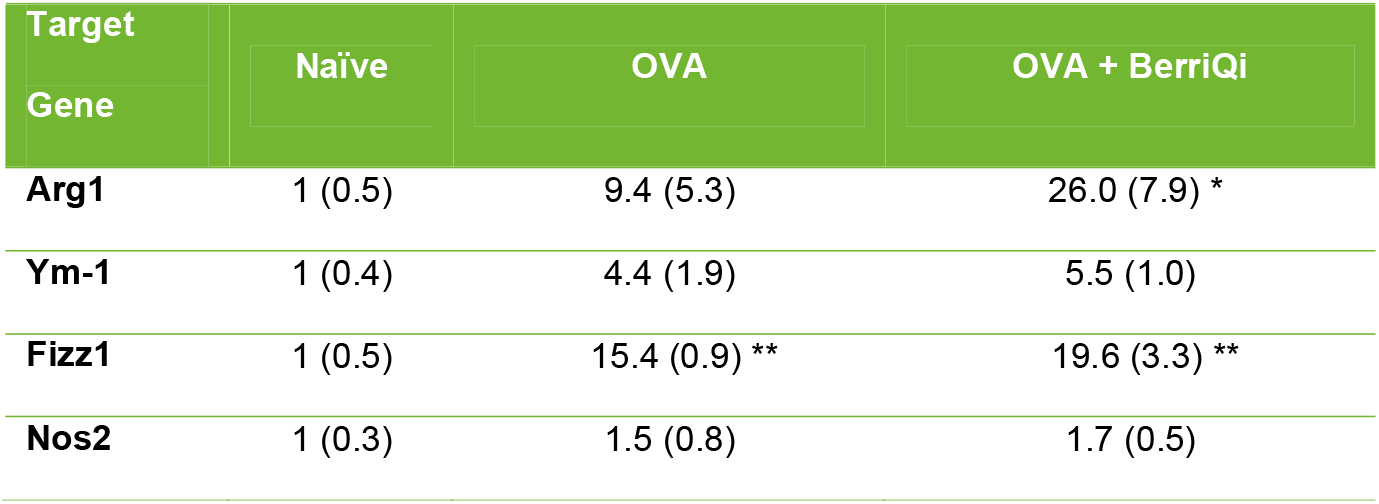
BerriQi^®^ Boysenberry and apple juice concentrate increases alternatively activated macrophage gene expression in the lung. Mice were primed with ovalbumin (OVA)/Alum i.p. and then challenged 7 days later with OVA i.n. (Day 0). Mice were orally gavaged with 2.5 mg/kg total anthocyanins (TAC) in the BerriQi^®^ Boysenberry and apple juice concentrate (BerriQi) 1h before OVA challenge and again 2 days post-challenge. Mean fold-change (SEM) in gene expression was measured by real-time qPCR in lung tissue 4 days post-OVA challenge. *P<0.05, **P<0.01 compared with naïve (one-way ANOVA with Tukey’s Post Hoc test) for 4 experimental replicates with n=3 per treatment groups.

| Target Gene | Naïve | OVA | OVA + BerriQi |
| --- | --- | --- | --- |
| **Arg1** | 1 (0.5) | 9.4 (5.3) | 26.0 (7.9) * |
| **Ym-1** | 1 (0.4) | 4.4 (1.9) | 5.5 (1.0) |
| **Fizz1** | 1 (0.5) | 15.4 (0.9) ** | 19.6 (3.3) ** |
| **Nos2** | 1 (0.3) | 1.5 (0.8) | 1.7 (0.5) |

Consumption of 2.5 mg/kg TAC BerriQi^®^ Boysenberry and apple juice concentrate led to increased levels of the cytokines IL-17A, CXCL10, and CCL4 (Figure 3B-D) 4 days following OVA challenge, but did not affect the IL-17F concentration (Figure 3E). We saw no effect on the concentrations of the classical Th1/2 cytokines IFNγ, IL-5, IL-9 or IL-10 in either the BerriQi^®^ Boysenberry and apple juice concentrate treated or the OVA alone mice compared to naïve controls (Figure 2C-F).

## 4. Discussion

We evaluated the effects of dietary supplementation with 2.5 mg/kg TAC BerriQi^®^ Boysenberry and apple juice concentrate, on the immune responses in a mouse model of acute allergic airways inflammation. Our results show that consumption of 2.5 mg/kg TAC BerriQi^®^ Boysenberry and apple juice concentrate reduced granulocyte and local T cell infiltration into the lung after OVA challenge, but did not alter T cell activation within the lung draining lymph node or the levels of classical Th-2 and Th-1 cytokines in the lung at four days following OVA challenge. Our current results indicated that BerriQi^®^ Boysenberry and apple juice concentrate had little impact on the Th-2/Th-1 mediated allergic response of mice, but rather targeted innate proinflammatory immune pathways. This is consistent with our previously reported finding in a mouse model of chronic allergic airways inflammation using 10 mg/kg TAC Boysenberry juice concentrate (33). Chemical composition analysis showed that the BerriQi^®^ Boysenberry and apple juice concentrate formulation contained high concentrations of cyanidin glycosides, ellagitannins, and chlorogenic acid. These compounds have been previously shown to reduce inflammatory signalling in vitro (38, 41, 42), and in vivo animal models of inflammation (34, 43–46). Our current results suggest that consumption of 2.5 mg/kg TAC BerriQi^®^ Boysenberry and apple juice concentrate, which also contains high levels of ellagitannins and chlorogenic acid, could have broader lung health benefits beyond allergic asthma disease by promoting the resolution of inflammation caused by innate immune cell overactivation. Current asthma therapies also suppress both the adaptive and innate immune responses without affecting the aberrant sensitivity to the allergen and this can lead to adverse events.

Consumption of BerriQi^®^ Boysenberry and apple juice concentrate had less of an effect on monocyte/macrophage infiltration into the lung than on granulocyte infiltration, and there was increased percentage of CD206+ monocytes. This could represent a shift to an M2 anti-inflammatory phenotype. We then measured the changes in gene expression for Arg1, Ym-1 and Fizz1, the classic genes for identifying alternatively activated macrophages (47, 48). Arg1 and Fizz1 gene expression was significantly increased with BerriQi^®^ Boysenberry and apple juice concentrate consumption. Consistent with a shift to a more anti-inflammatory macrophage phenotype, we detected increased levels of CXCL10 and CCL4 cytokines, which are produced by M2 macrophages, in the lungs of mice that consumed BerriQi^®^ Boysenberry and apple juice concentrate. These results suggest that the consumption of Boysenberry and apple juice concentrate led to a switch to the M2 phenotype in OVA-challenged mice. This could be one of the mechanisms by which BerriQi^®^ Boysenberry and apple juice concentrate consumption contributed to the resolution of inflammation. Previously, we reported that 10 mg/kg TAC Boysenberry juice concentrate can increase the abundance of alternatively activated (M2) macrophages, which promote tissue repair in a chronic model of airways inflammation (33). Our current results suggest that BerriQi^®^ Boysenberry and apple juice concentrate, had a similar effect in this mouse model of acute inflammation, in particular the increase in Arg1 gene expression is similar to our previously reported study showing increased arginase protein expression by alternatively activated macrophages (33). Further, research looking at an animal model Th2-mediated inflammation has identified M2 macrophage derived Fizz1 as a key limiting factor for Th2-mediated pulmonary inflammation (49).

The mice that consumed 2.5 mg/kg TAC BerriQi^®^ Boysenberry and apple juice concentrate showed increased levels of the cytokines IL-17A, CXCL10, and CCL4, but the levels of IL-17F were not affected. High IL-17 and IL-17F levels have been implicated in asthma pathogenesis, however, there is also evidence that IL-17A (50) can increase the abundance of MMP-9, an important tissue remodelling protein in asthma (33) as well as inducing apoptosis of neutrophils and eosinophils (50). CXCL10 and CCL4 are chemokines that attract monocytes/macrophages, and CXCL10 may also inhibit the infiltration of eosinophils in response to allergic airways inflammation (51). The combination of increased IL-17A-mediated granulocyte apoptosis and CXCL10-mediated inhibition of granulocyte infiltration could explain how the consumption of the BerriQi^®^ Boysenberry and apple juice concentrate resulted in decreased allergic airways inflammation, in particular the reduced number of eosinophils and neutrophils.

This switch to a M2 macrophage phenotype may be through the Boysenberry and apple polyphenols identified in the BerriQi^®^ Boysenberry and apple juice concentrate directly inhibiting proinflammatory pathways, or through an indirect shortening of the proinflammatory phase. Other studies have shown that increased dietary fibre shortened the duration of the proinflammatory phase, leading to reduced tissue damage in a chronic mouse model of allergic asthma (52). Based on the estimated fibre content in BerriQi^®^ Boysenberry and apple juice concentrate (Supplementary table 1), it is unlikely that the dietary fibre component played a significant role in the immune modulation we observed. However, the combination of anthocyanins with other polyphenols identified in the BerriQi^®^ Boysenberry and apple juice concentrate could have a similar effect on inflammation either by shortening the proinflammatory time course, or promoting the production of anti-inflammatory proteins. Ellagitannins have been shown in cell culture and animal models of chronic inflammatory diseases to reduce proinflammatory prostaglandins (53), cytokines (45, 54), and other proteins (42, 55, 56). Anthocyanins have also been shown to inhibit proinflammatory proteins (57, 58), and activate anti-inflammatory pathways in models of inflammation (59–63). It is possible that the combination of the different polyphenols in the BerriQi^®^ Boysenberry and apple juice concentrate act on a number of different immune pathways to regulate the immune responses to OVA.

We found that mice that consumed BerriQi^®^ Boysenberry and apple juice concentrate had reduced immune cell infiltration in response to acute OVA challenge and this could be as a result of a shift towards an anti-inflammatory environment within the lung. These results highlight the potential of anthocyanin-rich Boysenberry and apple dietary supplementation to modulate innate immune pathways during acute allergic lung inflammation. Further work is needed to determine if these pathways are also altered in other lung inflammatory conditions, such as air pollution exposure. Clinical studies are needed to show if these findings are translatable to human health.

## Author Contributions

OMS designed, performed, analyzed and interpreted the in vivo studies, and wrote and edited the manuscript; JC performed, analyzed and interpreted the chemcial composition experiments and GMS performed, analyzed and interpreted in vivo studies, and both contributed to the writing and editing of the manuscript; HD and SM performed the in vivo studies and helped edit the manuscript; RDH designed and directed the overall research programme and helped edit the manuscript.

## Acknowledgements

The authors acknowledge the efforts of TC Chadderton, and Brendan Vercoe in securing internal funding for this work. The authors would like to thank Andrew Carroll for assisting with data analysis and Jenny Smith for her invaluable help preparing the manuscript.

## SUPPLEMENTARY DATA

**Supplementary Figure 1.**
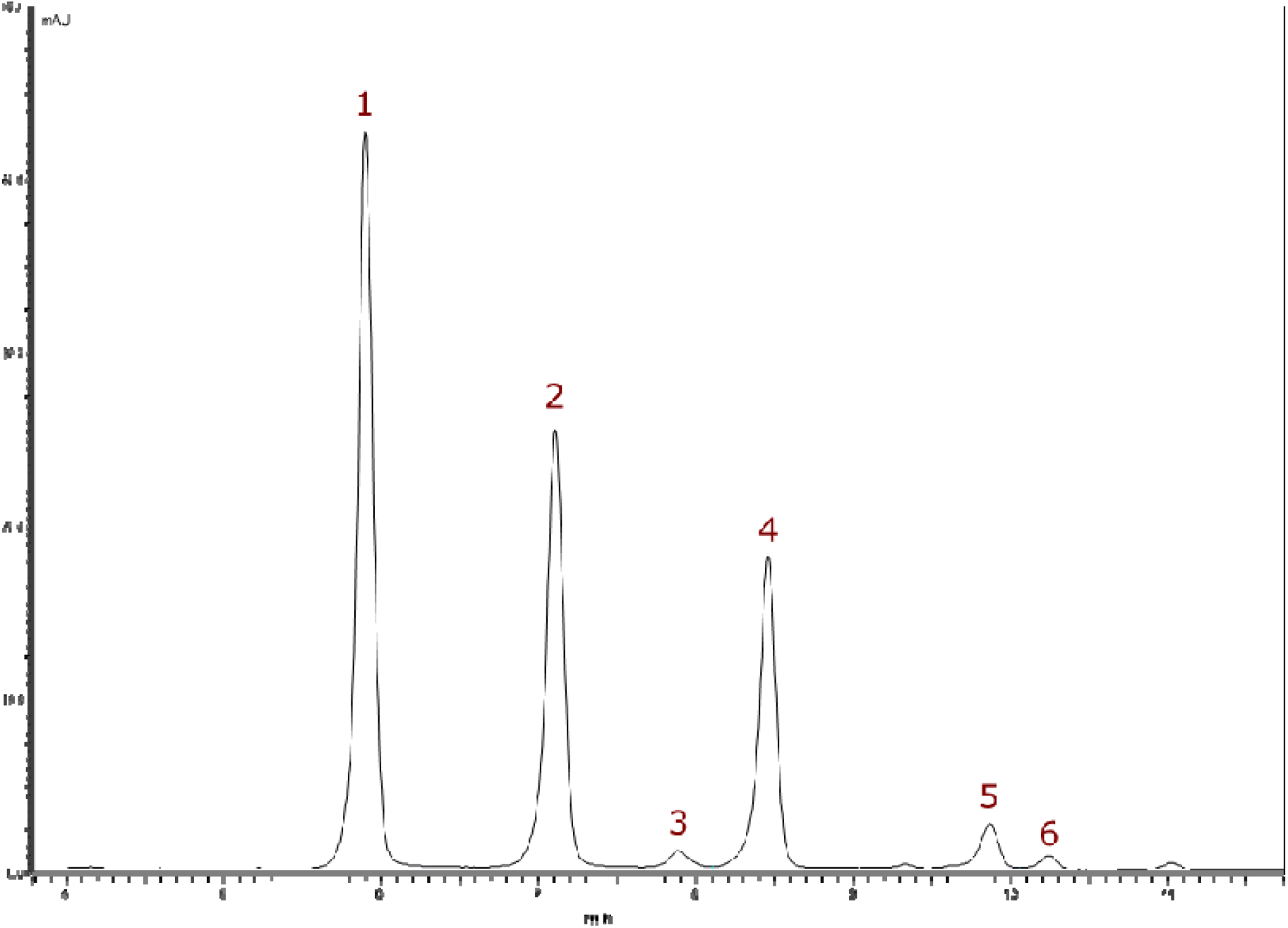
Liquid Chromatography-Mass Spectrometry (LC-MS) chromatogram for BerriQi^®^ concentrate, showing the anthocyanin UV/VIS profile measured at 520 nm. Peak numbers refer to compounds listed in Table 1.

**Supplementary Figure 2.**
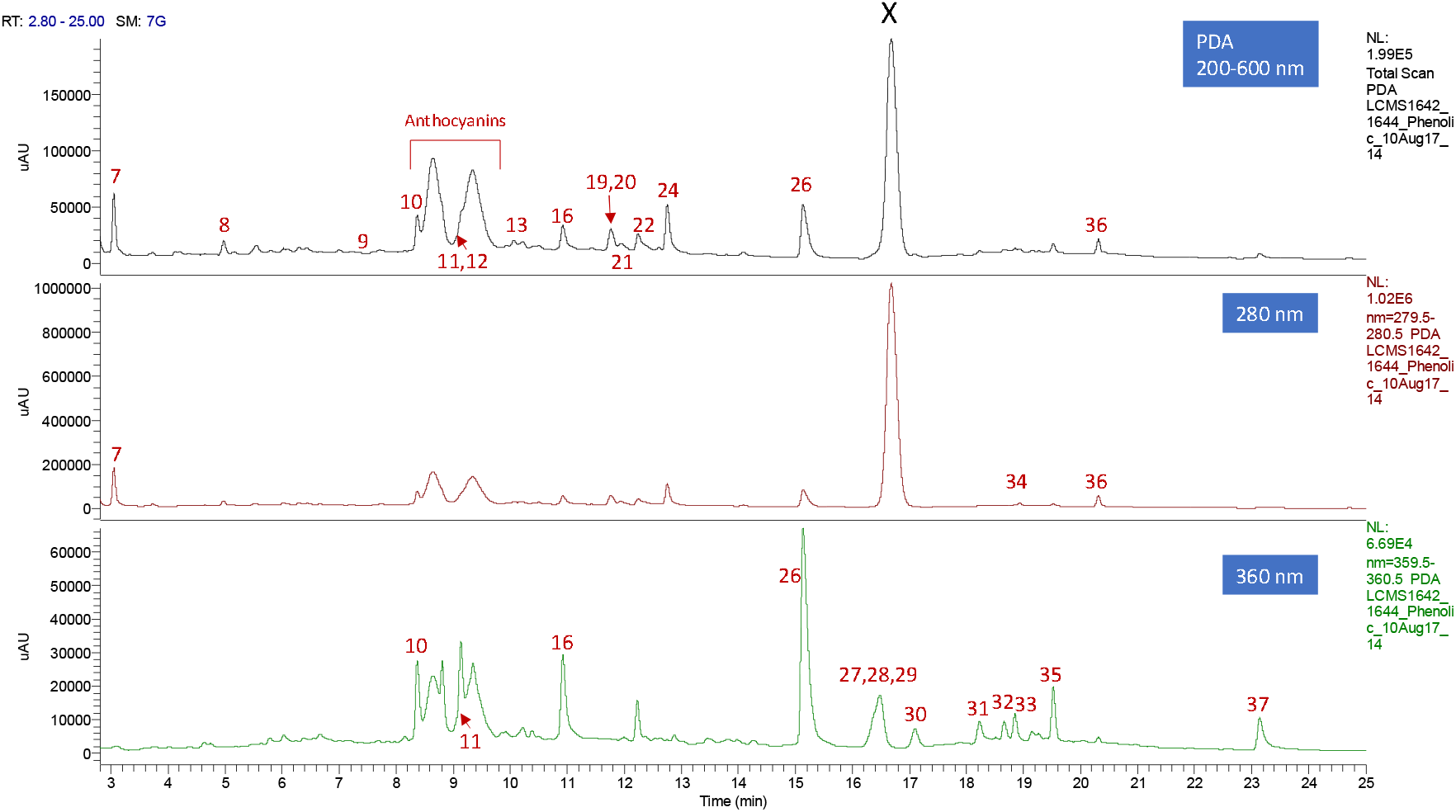
Liquid Chromatography-Mass Spectrometry (LC-MS) chromatograms for BerriQi^®^ concentrate, showing PDA (photodiode array) phenolic profiles measured at 200–600 nm, 280 nm and 360 nm. Peak numbers refer to compounds listed in Table 1. X, denotes sorbic acid, a preservative added to the BerriQi concentrate during formulation. This was present at a concentration of ~2000 μg/mL.

